# Oral microbiome and DNA virome profiling in pigs reveals age-associated dynamics and extracellular recovery of *Streptococcus suis-associated* phage lineages

**DOI:** 10.64898/2026.11.21.768310

**Authors:** Emmanuel Kuffour Osei, Mario Andre S. Ornelas, Edgar Garcia Manzanilla, Jennifer Mahony, John G. Kenny

**Affiliations:** School of Microbiology, University College Cork, Co. Cork, T12 K8AF, Ireland; APC Microbiome Ireland, University College Cork, Co. Cork, T12 YT20, Ireland; Food Bioscience, Teagasc Food Research Centre, Moorepark, Co. Cork, P61 C996, Ireland; Pig and Poultry Research and Knowledge Transfer Department, Teagasc Animal and Grassland Research and Innovation Centre, Moorepark, Fermoy, Cork, P61 C996, Ireland; School of Veterinary Medicine, University College Dublin, Co. Dublin, D04 V1W8 Ireland; Section for Food Microbiology, Gut Health and Fermentation, Department of Food Science, University of Copenhagen, Rolighedsvej 26, 1958 Frederiksberg C, Denmark; VistaMilk SFI Research Centre, Fermoy, Co. Cork, P61 C996, Ireland

## Abstract

The pig oral cavity is a critical interface for microbial transmission and herd health monitoring, yet its microbiome and virome remains poorly characterised. We performed shotgun metagenomic sequencing of oral fluids from pigs across 15 commercial farms in Ireland, using paired bulk and virus-like particles (VLP) enrichment to capture comprehensive microbial and DNA virome diversity. Microbial communities shifted progressively with age, with early post-weaning animals enriched in environmental genera such as *Pseudomonas* and *Staphylococcus* and later stages dominated by lactic acid bacteria and anaerobes. Only 20.6% of vOTUs could be assigned to a known genus, with the most abundant being *Astrithrvirus*, which increased with age. Both microbiome and DNA virome exhibited predictable, coordinated age-associated succession with strong cross-domain coupling supported by virus-host networks. Functional profiling of the microbiome revealed that predicted carbohydrate substrates tracked the dietary transitions, while accessory viral functions partitioned by lifestyle and host. The early post-weaning stage was characterised by high abundance of viral integration and excision functions, as well as elevated resistome diversity that was almost exclusively of prokaryotic origin rather than viral. Importantly, 26 of the 25,223 vOTUs identified were predicted to target *Streptococcus suis,* providing the first metagenomic catalogue of such phages from pig oral environments. These findings expand our understanding of oral microbiome-virome dynamics and establish a framework for integrated microbiome and viromics approaches that could enable therapeutic phage discovery and inform microbiome-targeted interventions in pig production systems.

## Introduction

Pigs represent one of the most economically significant livestock species, with over one billion animals raised annually to support global food security [1]. Beyond their agricultural importance, pigs serve as reservoirs for zoonotic pathogens, making microbial communities they harbour critically relevant to both animal and public health. The pig microbiome plays fundamental roles in nutrition, immune development, and colonisation resistance against pathogens [2]. However, when disrupted, these microbial communities can promote disease and pathogen emergence [3]. Consequently, understanding how the microbiome and virome develop across different pig production stages is crucial for both animal and public health.

The oral cavity occupies a pivotal position at the host-environment interface. While faeces index the distal gut microbiome, oral fluids integrate microbial signals from the mouth and the respiratory tract, as well as pen and feed-related environmental exposures. Due to this broad sampling capacity and non-invasive collection, oral fluids are already widely used in herd monitoring [4]. The swine gut microbiome and virome have received increasing attention in recent years. Faecal metagenomic data mining has identified tens of thousands of novel viral taxa, most with no significant hits to existing phage sequences in public databases [5]. Viruses are central to the ecology of microbial communities. Bacteriophages (or phages) in particular, shape the bacterial populations through predation, lysogeny, and horizontal gene transfer, which drives community turnover and evolution [6]. Many viruses encode genes that interact with host metabolism and cellular machinery. Auxiliary metabolic genes (AMGs)—are viral genes acquired from bacterial hosts that can augment or reprogramme host metabolic pathways to optimise conditions for viral production [7]. Phages also carry morons—accessory prophage genes that primarily benefit host fitness—as well as host-takeover proteins that subvert cellular machinery to suppress host defences and prioritise viral replication. Through these diverse mechanisms, viral infection can fundamentally alter bacterial metabolism and influence nutrient cycling [8]. In contrast to swine gut microbiome, the swine oral microbiome and virome remain poorly characterised. Recent studies have shown that oral bacterial communities are distinct from those in the swine gut and the housing environment, typically being less diverse but dominated by Bacillota (Firmicutes), particularly streptococci [9]. Earlier culture-based studies established foundational knowledge of oral pathogens such as *Streptococcus suis* and *Actinobacillus pleuropneumoniae* [10, 11], though such approaches are limited to cultivable taxa and cannot capture the full microbial diversity. However, the viral component of the oral cavity is virtually underexplored, despite the fact that oral viromes may act as reservoirs of phages that regulate bacteria, facilitate colonisation, and contribute to zoonotic spillover [12].

Methodological limitations in widely used viromics pipelines hinder virome research. Standard bulk metagenomic approaches, where viral sequences represent only a small fraction of total reads, typically recover abundant viruses while missing rare taxa [13]. Virus-like particle (VLP) enrichment can improve viral genome recovery by depleting host DNA, but this approach can be biased towards lytic forms and may underrepresent larger viruses [14, 15]. While each method introduces distinct biases, most swine virome studies have relied solely on bulk sequencing. Integrated approaches combining both bulk and VLP datasets remain limited, particularly for oral fluid.

Moreover, antimicrobial resistance (AMR) poses a major challenge in swine production systems where antibiotics are routinely used [16]. Pig microbiomes harbour extensive antibiotic resistance gene (ARG) repertoires, though the role of phages in ARG dissemination is a subject of ongoing debate [16–18]. Mining viromes can also reveal phage-encoded lysins and depolymerases with potential antibacterial activity [19]. As phage therapy gains renewed interest in both human and veterinary medicine, commercial phage products targeting pathogens, including *Salmonella*, *Escherichia coli*, and *Clostridioides* (*Clostridium)* species, are already available for applications in pig and other livestock [20]. However, several important pathogens remain poorly explored as potential targets for phage biocontrol. One such bacterium is *Streptococcus suis*. *S. suis* causes significant economic losses through meningitis and septicaemia in pigs while posing zoonotic risks to humans, particularly those in close contact with swine [21]. It is a genetically diverse species with several studies documenting the widespread occurrence of AMR genes and associated phenotypes [22]. Despite its importance, very few infective *S. suis* phages have been reported. Until recently, only a single infective phage had been isolated and genome-sequenced, with two additional phages subsequently reported in 2025 by our group [23, 24]. Also, our previous works have revealed extensive prophage load and anti-viral defence systems within *S. suis* genomes yet attempts to isolate extracellular phages from pig-derived samples that infect this bacterium have been largely unsuccessful [20, 25].

In this study, we address these knowledge gaps through the first comprehensive, combined metagenomic characterisation of the pig oral microbiome and DNA virome across different age groups. Using paired bulk and VLP sequencing of oral fluids collected from large commercial farms across Ireland, we demonstrate that prokaryotic and viral communities shift in parallel with age, with strong coupling between their diversity and compositional patterns. Functional analysis reveals abundant viral AMGs and host-takeover genes, while antimicrobial resistome profiling highlights distinct age-associated ARG distribution patterns. Importantly, we recover extracellular viral species predicted to infect *S. suis*, establishing a metagenomic catalogue of such viruses from pig oral fluids. These findings expand our understanding of the oral microbiome-virome ecosystem, provide a methodological framework for integrated viral characterisation, and reveal the presence of a novel phage pool with direct relevance to swine health management and zoonotic disease control.

## Materials and methods

### Ethics approval

The study protocols carried out in this work were reviewed and approved by the Teagasc Animal Ethics Committee (reference 0123-368). Teagasc is the Irish Agriculture and Food Development Authority and undertakes research, education, and dissemination of knowledge nationwide in the Republic of Ireland.

### Farms, sampling and processing

Oral fluid samples were collected from pigs on eighteen farrow-to-finish farms (farm size range 130 to 2,400 sows) across Ireland from March to July 2022, as previously described [4]. In brief, cotton ropes (Hipra, Amer, Spain) were suspended at shoulder height within pens to allow pigs to chew on them. To mitigate contamination, ropes were positioned away from pen walls, feeders, and drinkers or held by an operator when required. After chewing, ropes were transferred into a plastic bag fitted to a 10 ml tube, and oral fluids were recovered by squeezing the rope into the tube.

Sampling was carried out at the pen level at four production stages (age groups); one week after weaning (5 weeks old, Weaner 1), one week prior to transfer to the finishing facility (10– 11 weeks old, Weaner 2), one week after transfer to the finishing facility (12–13 weeks old, Finisher 1), and one week before slaughter (22–25 weeks old, Finisher 2). Four ropes were collected from four different pens at each stage, yielding 16 samples per farm. For the present study, oral fluids from fifteen of the eighteen farms originally visited were used, representing nine counties in the Republic of Ireland (Table S1). In total, 240 samples were available for DNA extractions (15 farms × 4 age groups × 4 pens).

### Bulk microbiome and VLP DNA extractions

Prior to microbiome DNA extractions from oral fluids, samples from the four pens within each age group on a given farm were pooled, resulting in 60 composite samples (15 farms × 4 age groups). DNA was extracted from aliquots of composite oral fluid samples using QIAamp PowerFecal Pro DNA Kit (Qiagen, Germany, #51804) according to manufacturer’s instructions. One negative control consisting of Milli-Q water, and one positive control consisting of the ZymoBIOMICS Oral Microbiome Standard (Zymo Research Inc., USA, #D6332) were processed using the same kit and procedure as for the microbiome samples.

For VLP DNA extractions, 4 ml aliquots of oral fluids were centrifuged at 5,000 ×*g* for 15 minutes. The supernatants were filtered through a 0.45 µm polyethersulfone membrane filter (Sarstedt) twice to remove cellular fraction. Extractions were carried out using the Norgen Phage DNA Isolation Kit (Norgen Biotek Corp., Canada, #46800) with some modifications to manufacturer’s instructions. Briefly, filtrates were treated with 20 U of DNase and 10X DNase buffer (Thermo Fisher Scientific, USA, #AM2238), and 1 µL of RNase A (Thermo Fisher Scientific, USA, #EN0531). The reaction was incubated at 37 °C for 70 minutes to digest non-encapsidated nucleic acids followed by nuclease inactivation with 20 mM EDTA. Four microlitres of proteinase K (20 mg/ml) and 500 µL of lysis buffer was added and vortexed for 10 seconds. This mixture was incubated in a water bath for 30 minutes at 56 °C and subsequently treated with 320 µL isopropanol and purified on a column according to the manufacturer’s instructions. One negative control consisting of Milli-Q water was also processed using the same method as for VLP samples.

### Metagenomic sequencing of microbiome and VLP DNA, reads processing, and metagenome assembly

All extracted DNA was quantified using 1X dsDNA HS Kit (Invitrogen, #Q33231). Oral microbial DNA and VLP DNA extracts were used for library preparation and sequencing (Novogene, U.K.). DNA was normalised and sheared by sonication, followed by end-repair, dA-tailing and adapter ligation. Quality of indexed libraries was assessed using Agilent Tape Station with qPCR quantification. Samples were pooled and sequenced using 2 × 150 nt paired-end sequencing chemistry on the Illumina NovaSeq X Plus platform. Trim Galore (v0.6.10), which incorporates FastQC (v0.12.1) and Cutadapt (v4.7), was used to assess the quality of raw paired-end reads, trim low-quality base calls, and remove adapter sequences (https://github.com/FelixKrueger/TrimGalore). To remove host sequences, trimmed reads were aligned to the pig (*Sus scrofa* Sscrofa11.1) and human (GRCh38, no-alt assembly) reference genomes (https://benlangmead.github.io/aws-indexes/bowtie) using Bowtie2 (v2.5.3) in the “*very-sensitive*” mode [26]. On average, 22.2 ± 0.2 and 23.4 ± 0.4 million cleaned paired-end reads were generated for bulk microbiome samples and VLP samples, respectively. The sequencing control using Zymo oral mock community successfully detected all 12 expected species with most abundances within reasonable range of theoretical values, demonstrating adequate taxonomic resolution. All negative controls failed library QC, and no sequencing data could be generated. In addition, one VLP library failed QC, resulting in a final dataset comprising 60 bulk microbiome metagenomes and 59 VLP metagenomes (Table S1). *De novo* assemblies were generated for each cleaned dataset (both microbiome and VLP) using SPAdes (v4.0.0) in *“-meta”* mode [27].

### Viral identification and clustering into vOTUs

Viral contigs were identified and processed using the Modular Viromics Pipeline (MVP v1.0) [28]. The cleaned reads (generated from host decontamination described above), and corresponding assemblies of both bulk microbiome and VLP metagenomes were provided as input for MVP workflow. In the workflow, viral sequences were predicted with geNomad (v1.5.2) using a minimum contig length of 5 kb. Quality of these predicted viral contigs were assessed with CheckV (v1.0.1) and candidate viral contigs were filtered to retain those with at least one predicted viral gene and a host-viral gene ratio ≤ 1. Contigs with ≥ 50% CheckV completeness including medium quality, high quality, and complete viral contigs were retained. Contigs below the 50% CheckV threshold were retained if they had a geNomad virus score ≥0.95, encoded ≥3 viral genes and ≥2 viral hallmark genes, and were assigned a taxonomy resolved at least to the realm level. Because completeness-based filtering alone disproportionately removed divergent lineages from under-represented ecosystems [29, 30] such as the pig oral cavity, this additional criterion was applied to retain potential novel viruses. Filtered contigs were dereplicated into species-level viral operational taxonomic units (vOTUs) using BLAST+ (v2.12.0) followed by average nucleotide identity-based clustering, with thresholds of ≥ 95% ANI across ≥ 85% alignment coverage. Representative sequences were selected for each vOTU, and Bowtie2 (v2.5.1) index was built for downstream read mapping. Cleaned reads were mapped to vOTUs and coverage/abundance metrics were calculated with CoverM (v0.6.1). vOTU abundance tables were generated, and abundance was normalised using RPKM (reads per kilobase per million mapped reads). For downstream analysis, only vOTUs with ≥ 75% horizontal coverage across their representative sequence were retained to define presence in a sample (Table S2).

### Functional characterisation of the microbiome and DNA virome

To characterise microbial function, protein-coding genes were predicted from assembled contigs using Prodigal (v2.6.3) in metagenomic mode (-*p meta*). Functional annotation of the predicted proteins was performed with eggNOG-mapper (v.2.1.12) using DIAMOND against the eggNOG database (emapperdb-5.0.2) [31]. Gene abundances were quantified by mapping paired-end reads to their corresponding assemblies using BWA-MEM (v2.2.1), at thresholds of ≥95% read identity and ≥75% aligned read length [32]. Gene-level annotations were merged with abundance estimates and aggregated to KEGG Orthology (KO) and KEGG pathways. Carbohydrate-active enzymes (CAZymes) were annotated with dbCAN (v5.2.8) using the “*easy_substrate*” workflow [33]. Reads were mapped to assemblies using BWA-MEM (v2.2.1) and samtools (v1.20.0), requiring a mapping quality of ≥30, sequence identity of ≥0.98, and an overlap ratio of ≥0.2. Next, we generated TPM-normalised abundances for CAZyme families, predicted family substrates and carbohydrate gene clusters, including polysaccharide utilisation loci (PUL)-homology substrate predictions.

For viral genome annotation, Pharokka (v1.10.1) was used in metagenomic mode (--*meta*) [34]. Genes were predicted using *prodigal-gv*, and predicted proteins were assigned to PHROGs using MMseq2 in reverse search mode (--*reverse_mmseqs2*). To enhance the rate of annotation, genbank files from Pharokka were used as input for Phold (v1.2.6), which applies foldseek and colabfold for structural homology predictions [35]. For viral taxonomic assignment, host prediction, and lifestyle annotation, we used PhaGCN, CHERRY, and PhaTYP within PhaBOX2 (v2.1.13), respectively using “*--task end_to_end*” and “--*skip Y*” parameters (Table S3) [36]. Host predictions were integrated with abundance-weighted viral functional profiles.

### Antimicrobial resistome analysis

Antimicrobial resistome was characterised using both read-based and assembly-based approaches. For read-based approach, metagenomic reads from bulk microbiome were screened with resistance gene identifier (RGI v6.0.3) in *“bwt”* mode against the Comprehensive Antimicrobial Resistance Database (CARD v4.0.2), including variant models [37]. ARG abundance profiles were normalised for downstream analyses. For assembly-based approach, metagenomic assemblies from both bulk microbiome and VLP were independently annotated with RGI using “main” mode and DIAMOND as the alignment tool. Only “perfect” or “strict” hits were included in downstream analyses. To predict the microbial source of ARG-containing contigs, bulk microbiome assemblies were classified with Kraken2 [38]. RGI hits were mapped to Kraken2 contig-level taxonomic assignments using genomic coordinates. The taxonomic lineages of contigs carrying ARGs were then resolved using TaxonKit v0.16.0 [39]. This approach enabled the mapping of ARGs, resistance mechanisms, and drug classes to putative bacterial hosts where sufficient taxonomic information could be obtained from contigs.

### Taxonomic assignment and differential abundance analyses

Bulk microbiome reads were assigned taxonomic classifications using Kraken2 (v2.1.3) with the PlusPF database (https://benlangmead.github.io/aws-indexes/k2). To refine abundance estimates, Bracken (v2.9) [40] was then applied across taxonomic ranks, and outputs were merged across samples for downstream analysis. To reduce sparsity and focus on biologically relevant taxa, a 10% prevalence threshold was applied to filter out transient contaminants. Viral taxonomic assignments were obtained from PhaBOX2 as described above. Differential abundance of viral and microbial taxa was determined using ANCOM-BC2 (v2.8.1) in R [41]. Both microbiome and virome analyses incorporated age group and farm as covariates as well as accounting for extraction method (bulk or VLP) specifically for the virome analysis. Both pairwise comparisons and one-vs-rest analyses, in which each age group was contrasted with all other groups combined, were performed. Effect sizes from ANCOM-BC2 were reported as log-fold-change coefficients (β), with multiple-testing correction using the Benjamini-Hochberg correction procedure.

### Statistics and ecological measures

Analyses were performed in R (v4.3) using vegan (v2.6), lmerTest (v3.1.3), emmeans (v1.11.22), ape (v5.8), and phyloseq (v1.0). All R packages are available at https://cran.r-project.org/web/packages/. For the bulk microbiome, species-level Bracken profiles were filtered to include only taxa present in ≥10% of samples. Observed richness, Shannon diversity, and Simpson diversity were calculated, and the differences among groups were assessed using linear mixed-effects models with age group as fixed effect and farm as random intercept. Pairwise contrasts were obtained from estimated marginal means, with false discovery rate (FDR) correction applied. Beta diversity was assessed using Aitchison distance on centred log-ratio (CLR)-transformed species-level abundance data and visualised using principal component analysis (PCA) plots. Differences in community composition among age groups were tested using PERMANOVA with 999 permutations blocked within farm. To further account for farm-associated variation, distance based redundancy analysis (dbRDA) was performed as ∼ AgeGroup + Condition(Farm_ID), thereby isolating the effect of age group on community composition.

For the virome, RPKM values were set to zero for vOTUs where <75% of the vOTU genome was covered by mapped reads. Alpha diversity was characterised using observed vOTUs and Shannon diversity. Mixed effect models included age group, extraction method and their interaction as fixed effects, while farm and matched sample pair (bulk and corresponding VLP) were included as random intercepts to account for repeated measures and potential farm-level clustering. Pairwise contrasts between age groups were evaluated within each extraction method to assess potential methodological influences. Beta diversity was assessed using Bray-Curtis dissimilarities, and the effects of age group, extraction method, and farm were assessed using marginal PERMANOVA, with permutations constrained within farm. Additionally, dbDRA was used to examine the effects of age and extraction method after conditioning on farm.

Associations between microbial and viral communities were investigated using matched biological samples to enable direct comparison of diversity and composition. Concordance in alpha diversity was assessed for richness and Shannon indices using Spearman correlations. Beta diversity concordance was evaluated with Bray-Curtis, applying Mantel tests and Procrustes analysis with 9,999 permutations constrained within farm. Predicted virus-host relationships were further explored by correlating the abundance of bacteria host genera with the abundance of vOTUs assigned to the same host genus, using Spearman correlations and FDR correction.

Boxplots show median and interquartile range, with whiskers extending to 1.5 × IQR. Ordination plots include 95% confidence ellipses. Statistical significance is denoted as **** *p* < 0.0001, *** *p* < 0.001, ** *p* < 0.01, or *p* < 0.05.

### Detection of *Streptococcus suis*-targeting phages

Putative *S. suis*-associated vOTUs were identified using complementary host-prediction and sequence-homology methods. Initially, CHERRY host predictions generated within PhaBOX2 were screened to identify VOTUs assigned to *S. suis*. In parallel, we downloaded all phage genomes available as part of the INPHARED database (accessed on 16^th^ October 2026) [42]. The nucleotide sequences of all phages that infect bacteria within the *Streptococcus* genus were extracted and deduplicated using accession and phage names. Where duplicate sequences were identified, the RefSeq version of the pair was retained for downstream analysis. In total, 82 genomes of phages (including prophages) predicted to infect *S. suis* were retrieved. The 82 sequences were searched against all vOTUs using BLAST (v2.14.1). BLAST alignments were summarised across non-overlapping high-scoring pairs, and candidate vOTUs were retained if ≥3,000bp were aligned, ≥50% of the shorter sequence was covered, and the best HSP had ≥85% nucleotide identity. These thresholds were selected to facilitate candidate discovery and do not represent taxonomic boundaries. Candidates identified by either approach were merged and deduplicated to generate a non-redundant set (Table S4). To increase confidence in viral origin, Phold outputs were inspected to select vOTUs that were ≥10 kb in length and encoded ≥2 genes in “head and packaging” and “tail” functional category. Relative abundance profiles were extracted from virome abundance tables. Prevalence was defined at biological sample level, such that a vOTU was considered present if detected in either bulk microbiome or the corresponding VLP fraction or both, and expressed as the proportion of biological samples in which it occurred.

For comparative phylogenomic analysis, the candidate *S. suis* vOTUs and a non-redundant set of all streptococcal phages from INPHARED database were used as input for ViPTreeGen (v1.1.3) with default parameters (Table S5) [43]. The resulting tree was visualised in R using ggtree (v3.17.0). Pairwise intergenomic similarity among the candidate *S. suis* vOTUs and the 82 INPHARED *S. suis* phage sequences were estimated using VIRIDIC (v1.1) with default parameters, applying 70% and 95% thresholds for genus and species delineation, respectively [44]. The resulting similarity matrix and cluster tables were subsequently used to generate a hierarchical clustering heatmap.

## Results

### Oral microbiome composition and diversity shift across pig age groups

We analysed the total microbiome composition in 60 composite (pooled from 240 pen-level samples) oral fluid metagenomes collected cross-sectionally from four production stages (age group) across 15 farms. Observed species richness increased significantly following the earliest post-weaning stage, with clear differences among age groups (linear mixed model: *F* (3,42) = 9.18, *p* < 0.001) (Fig. 1A). Specifically, Weaner 1 had significantly fewer detected species compared to all other age groups (*p* < 0.001). This pattern in observed richness was consistent at the genus level (*F* (3,42) = 7.190, *p* < 0.001). In contrast, Shannon and Simpson diversity indices showed no significant differences between groups, indicating gains in the number of taxa rather than systematic shifts in evenness. Beta diversity analysis revealed a clear separation of microbial communities according to age groups. PCA of CLR-transformed abundance data separated Weaner 1 from older stages at both genus and species (Fig. 1B) levels. A PERMANOVA controlling for farm effects confirmed significant compositional differences among age groups (genus: *R^2^*=0.197, *p* < 0.001; species: *R^2^*=0.147, *p* < 0.001). While the earliest age group showed greater within-group dispersion, the age effect remained robust under blocked permutation testing. Distance-based redundancy analysis (dbRDA) using Bray-Curtis dissimilarity and conditioning on farm effects demonstrated consistent age-based separation, supporting the observation that compositional turnover follows a developmental trajectory that is independent of farm-specific factors (Fig. S1A).

**Fig. 1:**
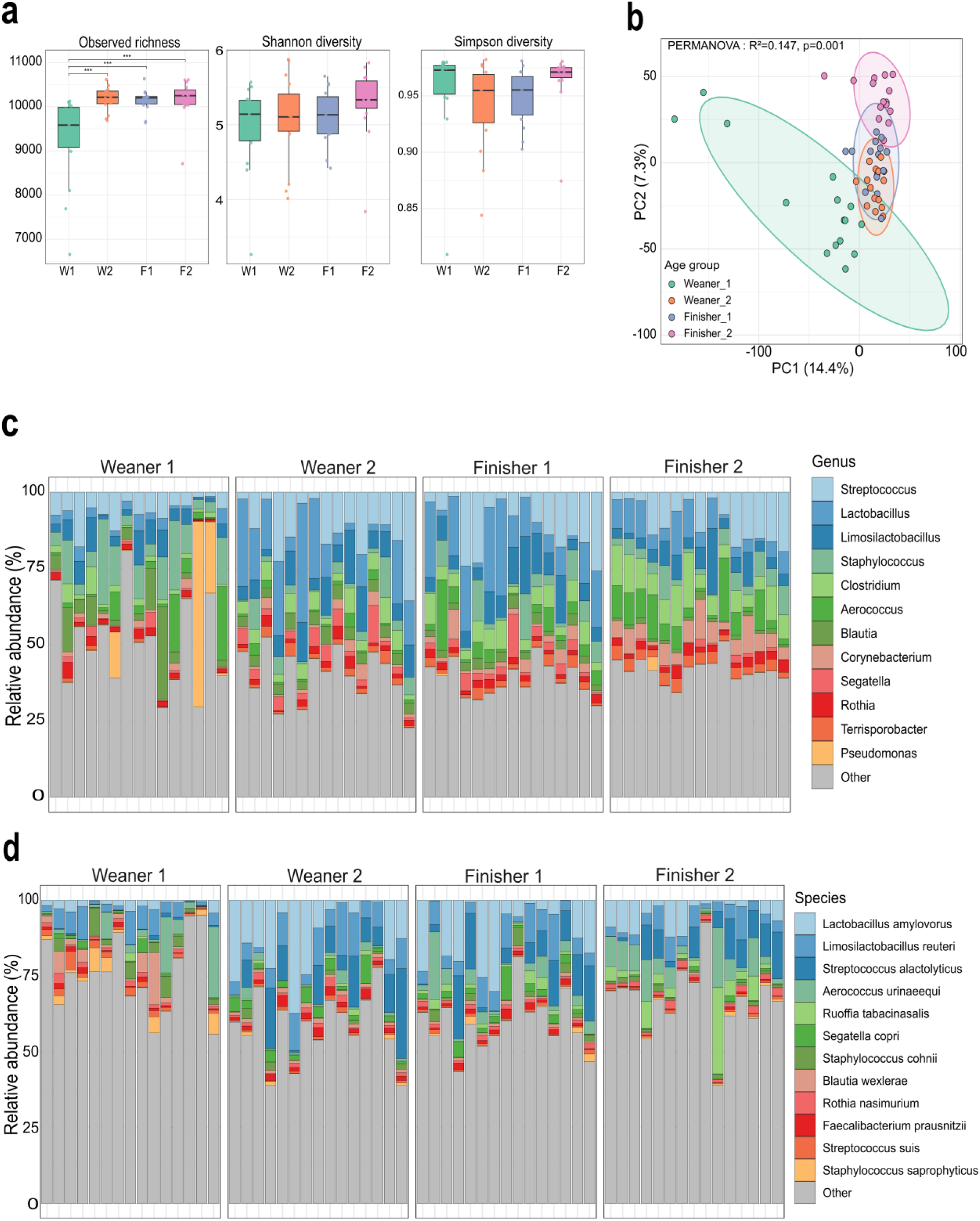
Changes in oral microbiome diversity and composition across age groups. **(a)** Alpha diversity metrics showing observed species richness (left), Shannon diversity (middle), and Simpson diversity (right) across age groups. Observed species richness increased significantly after earliest post-weaning stage, with Weaner 1 having fewer species than all other groups (FDR-adjusted pairwise q < 0.001). W1, W2, F1, and F2 represents Weaner 1, Weaner 2, Finisher 1, and Finisher 2, respectively. **(b)** Principal component analysis ordination of oral microbiome species beta diversity measured by Aitchison distance with CLR-transformed abundance data. PCA separated Weaner 1 from older stages, with PERMANOVA confirming significant compositional differences among age groups (species: R^2^=0.147, p < 0.001). Points are coloured by age group. **(c)** Taxonomic composition at genus level showing relative abundance across individual samples within each age group. **(d)** Taxonomic composition at species level.

Microbial composition varied across age groups, reflecting the age-associated diversity patterns. At the genus level, Weaner 1 communities were characterised by relatively high average abundances of *Staphylococcus* (10.3%) and *Pseudomonas* (6.8%), along with *Streptococcus* (6.6%), *Blautia* (6.3%), and *Limosilactobacillus* (5.2%) (Fig. 1C). In Weaner 2 and Finisher 1, *Lactobacillus* (14.9% and 13.6%) and *Streptococcus* (12.8% and 11.5%) dominated with sustained contributions from *Limosilactobacillus* (8.5% and 8.3%) and *Clostridium* (4.9% and 5.6%). The last age group, Finisher 2, showed a more balanced composition with elevated levels of *Streptococcus* (9.9%), *Clostridium* (8.8%), *Aerococcus* (7.4%), *Lactobacillus* (7.1%), and *Corynebacterium* (6.2%). Despite these age-specific differences, *Streptococcus* (10.2%), *Lactobacillus* (9.5%), and *Limosilactobacillus* (7.0%) remained consistently abundant across all groups, with a long tail of low-abundance taxa. At species resolution, Weaner 2 and Finisher 1 were characterised by *Lactobacillus amylovorus* (13.0% and 11.1%) and *Streptococcus alactolyticus* (9.4% and 8.0%) (Fig. 1D). *Limosilactobacillus reuteri* maintained a consistent presence across all stages while Finisher 2 showed elevated levels of *Aerococcus urinaeequi* (6.9%) and *Ruoffia tabacinasalis* (4.9%).

### VLP enrichment expands species recovery and reveals an oral virome composition structured by age

We next resolved the oral virome using VLP libraries alongside bulk metagenomes (60 bulk, 59 VLP after one VLP failed QC). Initial viral sequence identification with geNomad yielded 189,938 candidate viral contigs, including 4,365 putative prophages in bulk extraction. Following viral contig QC described above, we retained 67,777 viral contigs for downstream analyses. Of the 67,777 viral sequences, 64,375 (94.9%) represented MIUVIG genome fragments and 3,402 (5.01%) were classified as high quality (Fig. 2A). The retained contigs were predominantly assigned as prokaryote-infecting (99.6%) dsDNA viruses (99.8%), and lifestyle annotations indicated slightly more virulent viruses (51.7%) than temperate (42.9%) counterparts. VLP enrichment contributed to 88.3% of the 67,777 retained viral sequences. Clustering the viral candidates produced 25,223 species-level vOTUs (including 247 with prophages as representatives). VLP enrichment yielded a higher recovery of vOTUs, with 24,683 (97.9%) vOTUs detected, compared to 11,686 (46.3%) recovered from bulk fraction (Fig. 2B). Of all vOTUs, 11,146 (44.2%) were shared between both approaches, while 13,537 (53.7%) were unique to the VLP-enriched fraction and 540 (2.1%) unique to bulk fraction. This pattern was consistent within matched samples, where the median number of vOTUs recovered following VLP enrichment was 5,499, compared to 926 from bulk fraction (*p* < 0.0001; Fig. 2C). An age-stratified analysis indicated that the largest intersections of vOTUs extended across multiple age groups, with smaller age-specific viral populations, consistent with progressive turnover across the nursery-to-finisher transition (Fig. 2B). To assess whether the observed viral diversity approached saturation or whether additional sampling would be expected to recover more vOTUs, we examined viral richness as a function of sampling effort (Fig. 2E). In the combined dataset, a power-law accumulation model provided a better fit than an asymptotic exponential model (ΔAIC=84.0), with an estimated mean increment of approximately 69 vOTUs at the final sampling step. Similar results were observed for both the bulk (ΔAIC=129.3) and VLP-enriched ΔAIC=85.3) fractions. These results suggest that further sampling would likely yield additional viral diversity.

**Fig. 2:**
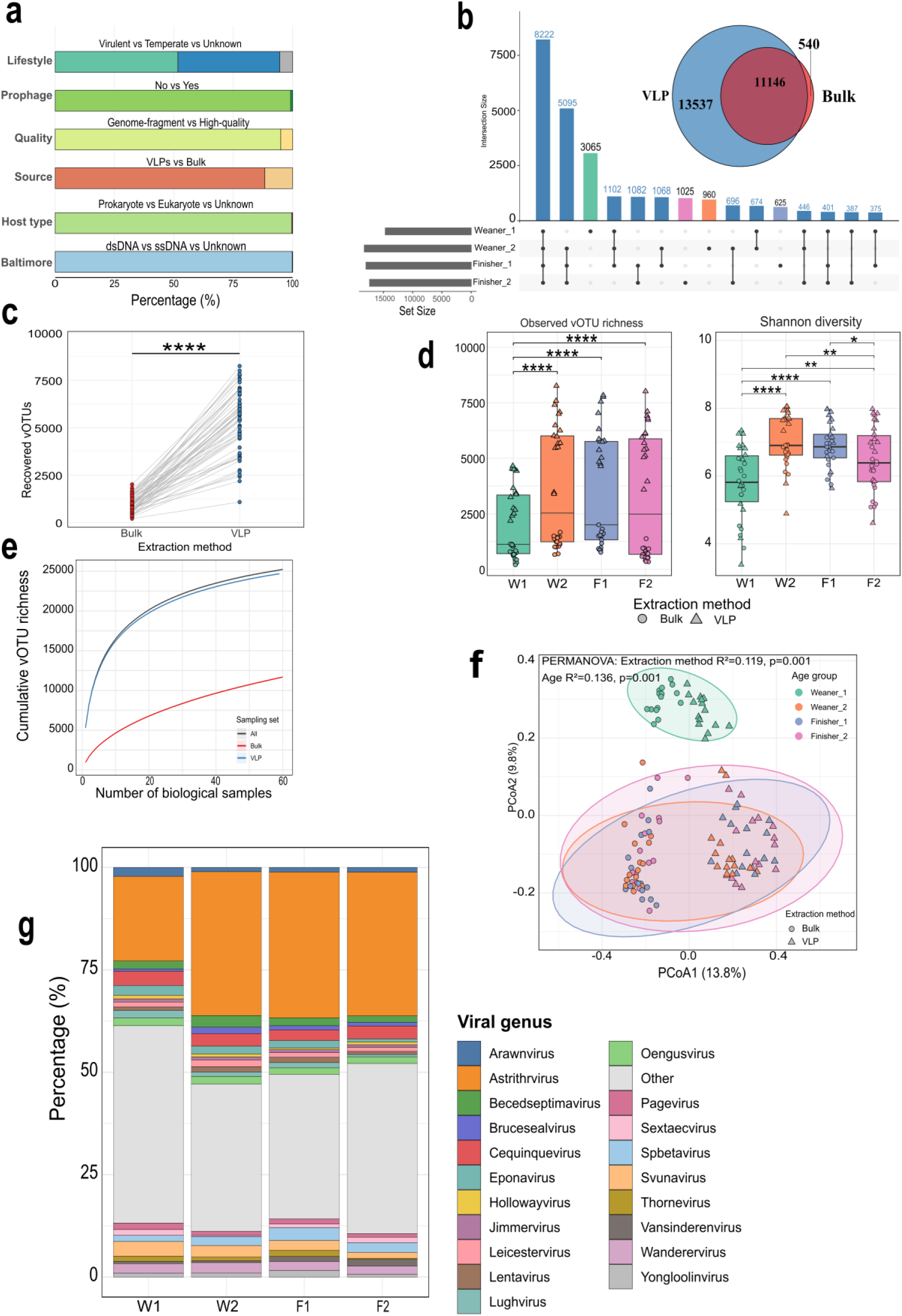
Viral community composition, recovery, diversity, and structure across pig age groups. **(a)** Genome characteristics of 67,777 retained viral contigs. Stacked bars show proportions classified by lifestyle (virulent, temperate, unknown), prophage, MIUViG quality, host type, and Baltimore class. Colours/stacks are ordered according to the labels shown on each bar. **(b)** Venn diagram showing vOTU (25,223) detection overlap between bulk and VLP extraction methods (top) and UpSet plot displaying age-stratified vOTU intersections (bottom). **(c)** Paired comparison of vOTU recovery from matched bulk and VLP libraries, showing significantly greater recovery following VLP enrichment (p < 0.0001). **(d)** Alpha diversity across age groups, showing observed vOTU richness (left) and Shannon diversity (right). Both metrics were lowest in Weaner 1. **(e)** vOTU accumulation curves for combined, bulk, and VLP datasets showing continued increases in cumulative vOTU richness. **(f)** Principal coordinate analysis of viral community composition based on Bray-Curtis dissimilarity. PERMANOVA identified age (R² = 0.136, p = 0.001) and extraction method (R² = 0.119, p = 0.001) effects on virome composition. Points are coloured by age and shaped by extraction method with 95% confidence ellipses. **(g)** Relative abundance of the classified viral genera across age groups.

Virome alpha diversity varied substantially with age group (Fig. 2D). Weaner 1 exhibited significantly lower vOTU richness compared to all other groups (*p* < 0.0001). Shannon diversity followed a similar trend with Weaner 1 showing lower diversity than all groups (all *p* ≤ 0.01). Additionally, both Weaner 2 (*p* < 0.01) and Finisher 1 (*p* < 0.05) had significantly higher Shannon diversity than Finisher 2, suggesting a modest decline at the final production stage. Both age group and extraction method influenced community composition (Fig. 2F). Bray-Curtis PERMANOVA revealed significant independent effects of age group (R*^2^*=0.136, *p* < 0.001) and extraction method (R*^2^*=0.119, *p* < 0.001). A farm-conditioned dbRDA confirmed a significant age effect (both global and age term *p* < 0.001), reinforcing that virome composition varies by age independently of between-farm differences (Fig. S1B).

For taxonomic classification, 24,480 (97.1%) could be placed in a taxon at the class rank. The viral community was overwhelmingly dominated by *Caudoviricetes* (n=24,428; 96.8%). Other classes comprised ≤ 0.3% of total vOTUs including, *Megaviricetes, Tectiliviricetes, Faserviricetes, Malgrandaviricetes, Papovaviricetes, Naldaviricetes, Revtraviricetes, Arfiviricetes,* and *Herviviricetes.* At the family level, *Salasmaviridae* was the most abundant assigned family and increased from 15.3% in Weaner 1 to 28.1% in Finisher 2. Other abundant families include *Autographiviridae, Steigviridae, Herelleviridae, Suoliviridae,* and *Rountreeviridae* (Fig. S2). At the genus level, only 5,200 (20.6%) vOTUs could be assigned to 464 known viral genera, while majority remained unclassified. Within the unclassified set, 10,089 (40.0% of all vOTUs) were singletons, likely constituting novel, undescribed lineages missing in current databases. Among those classified, *Astrithrvirus* (1,584 vOTUs) was the most abundant genus across all age groups, increasing from 20.7% in Weaner 1 to 35% in later groups (Fig. 2G). Other genera detected include *Cequinquevirus* (162 vOTUs), *Svunavirus* (141), *Wanderervirus* (120), *Spbetavirus* (115), and *Becedseptimavirus* (110). All other identified genera were represented by <100 vOTUs.

### Distinct bacterial and viral signatures characterise early and late production stages

We tested age-associated difference using ANCOM-BC2, which indicated clear patterns associated with age in both microbiome and virome. At the species level of the microbiome, one-vs-rest analysis identified substantial number of differentially abundant taxa in Weaner 1 (2,583), Weaner 2 (2,764), and Finisher 2 (4,430), whereas only 19 taxa were detected in Finisher 1. Weaner 1 was marked by enrichment of *Pseudomonas*, particularly *Pseudomonas* sp. B21128 (β=5.06, *q*=1.63×10^-7^), which is consistent with the observed high relative abundance of *Pseudomonas* at this timepoint (Fig. 3A). In Weaner 2, enrichment of taxa such as *Dialister, Megasphaera, Segatella*, and *Faecalibacterium* was observed. The Finisher 2 stage exhibited the strongest late-stage signature, with increased abundance of *Nosocomiicoccus ampullae* (β=4.16, *q*=4.11×10^-10^), *Corynebacterioum frankenforstense* (β=3.75, *q*=3.30×10^-10^), and several *Anaerococcus* species. At the genus level, *Nosocomiicoccus, Ruoffia, Anaerococcus, Corynebacterium*, and *Turicibacter* were identified as prominent taxa associated with Finisher 2.

**Fig. 3:**
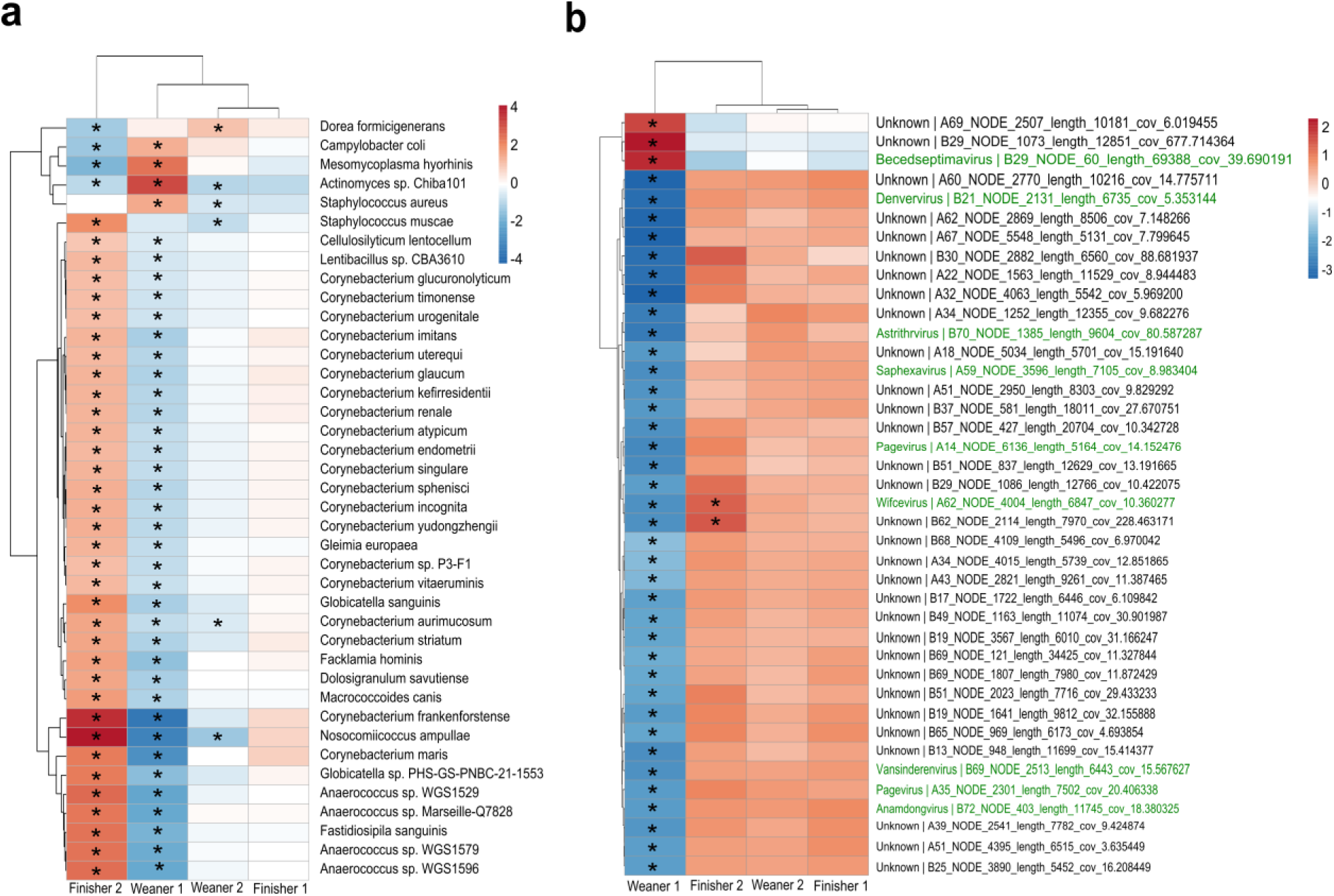
Differentially abundant prokaryotic species and viral populations across age groups. **(a)** Heatmap showing top 40 differentially abundant prokaryotic species and **(b)** Viral species differential abundance. Colours indicate ANCOM-BC2 log fold-change coefficients (β), with positive (red) values indicating enrichment and negative (blue) values indicating depletion in the focal group. Rows (taxa) and columns (contrasts) are hierarchically clustered based on β values. Asterisks (*) indicate significant differences at q≤0.05 with FDR correction.

Age-associated changes in the virome were most concentrated at the earliest and latest stages. One-vs-rest analysis identified 642 significant vOTUs in Weaner 1 of which 168 were enriched and 474 depleted, and 119 signficant vOTUs in Finisher 2, with 62 enriched and 57 depleted. No significant vOTUs were detected for Weaner 2 or Finisher 1. Similarly, pairwise comparisons indicated the greatest divergence in viral community composition between Weaner 1 and Finisher 2 (612 vOTUs), while Weaner 2 versus Finisher 1 did not show significant differences. Although majority of these significant taxa could not be assigned to a known genus, Weaner 1 was characterised by depletion of *Pagevirus, Spbetavirus*, and *Fischettivirus* vOTUs, whereas several *Astrithrvirus* vOTUs were enriched in Finisher 2 (Fig. 3B). These results indicate that the early weaning and late finishing stages are the most taxonomically distinct periods in the succession of oral microbial and viral communities.

### Prokaryotic and viral community structures are tightly coupled across the pig oral microbiome

Prokaryotic and viral community diversity showed coordinated variation across matched oral samples. A positive correlation was observed between prokaryotic richness and vOTU richness (Spearman ρ = 0.525, *p* < 0.0001; Fig. 4A), suggesting that samples with higher prokaryotic diversity also tended to harbour richer viral communities. This pattern was also evident at the level of whole community structure in which Bray-Curtus dissimilarities between the prokaryotes and virome were strongly correlated (Mantel r = 0.702, *p* = 0.0001; Fig. 4B). Furthermore, Procrustes analysis demonstrated close correspondence between their ordination configurations (correlation = 0.808, M^2^ = 0.347, *p* = 0.0001; Fig. 4C). Therefore, the similarity in prokaryotic community composition between pigs was generally accompanied by similarity in their viral assemblages, indicating coordinated structuring of the two domains. To investigate the ecological relationships that may contribute to the observe cross-domain concordance, we examined predicted virus-host associations. An abundance-weighted network comprising resolved prokaryotic virus-host associations revealed links involving several prominent oral taxa and multiple viral lineages (Fig. 4D). The strongest association was observed between *Salmonella* and *Astrithrvirus*. Other significant associations included *Lactobacillus-Cequinquevirus, Bacillus-Spbetavirus, Streptococcus-Fischettivirus*, and *Streptococcus-Vansinderenvirus*. Additional links were identified between *Clostridium* (*Clostridioides*) and *Brucesealvirus, Enterococcus* and *Saphexavirus, Faecalibacterium* and *Oengusvirus* or *Lughvirus, Prevotella* and *Burzaovirus, Nonlabens* and *Inhavirus*, and *Myxococcus* and *Myxoctovirus*. Several bacterial genera, such as *Aerococcus*, were also connected to putative novel viral genus clusters, while *Streptococcus, Carnobacterium*, and *Lactobacillus* were associated with both classified and putatively novel viral lineages.

**Fig. 4:**
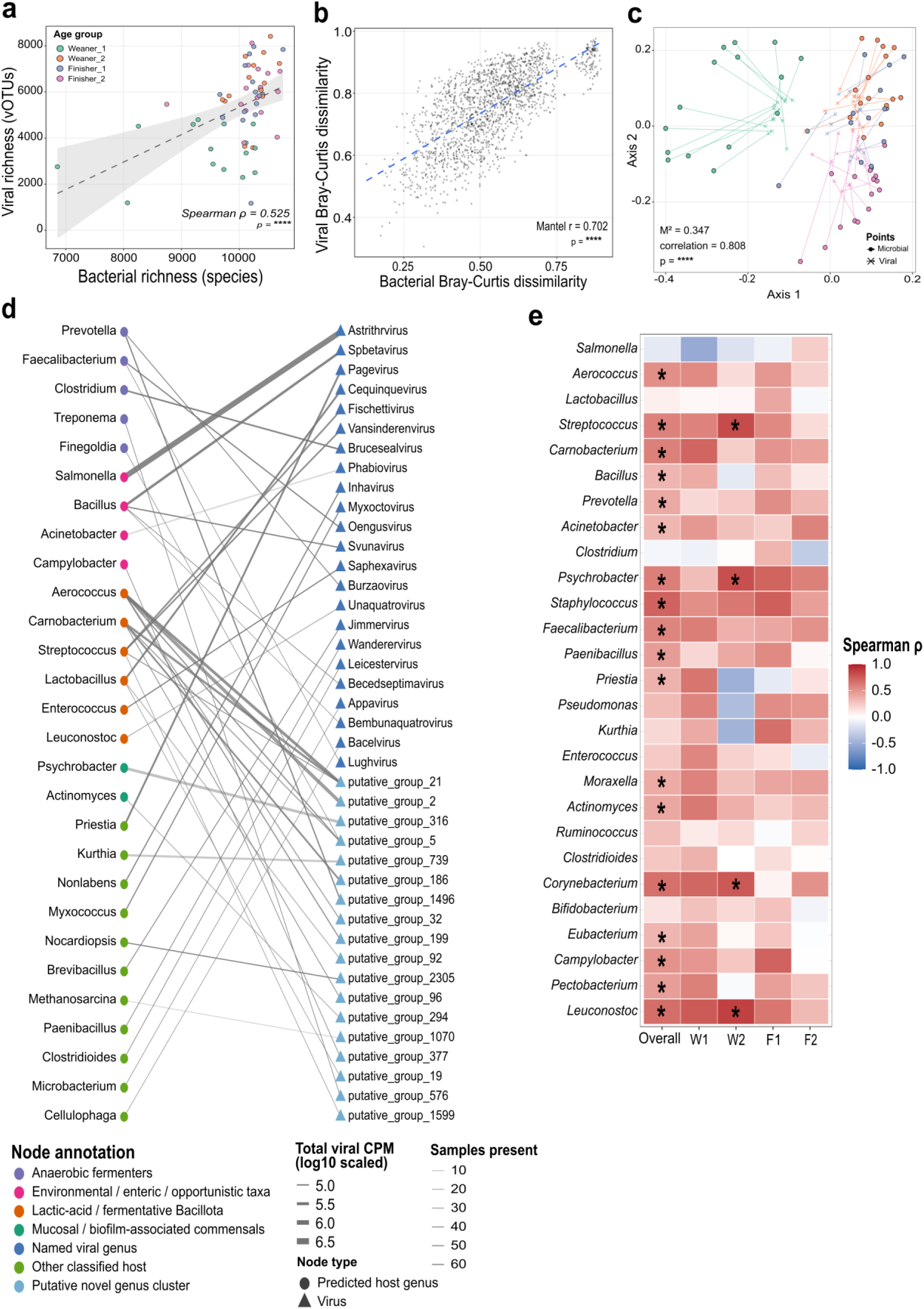
Coordination between oral prokaryotic and viral communities. **(a)** Association between prokaryotic species richness and vOTU richness across samples. **(b)** Mantel test scatterplot showing strong correlation between Bray-Curtis dissimilarities of prokaryotic and viral communities (Mantel r = 0.702, p < 0.001). **(c)** Procrustes overlay of PCoA ordinations confirming significant congruence between prokaryotic and viral community structures. For each sample, the prokaryotic ordination position is linked to the corresponding rotated viral position. Shorter connecting segments indicate greater agreement between the two community structures. The overall concordance was strong (M^2^ = 0.347, correlation = 0.808, p < 0.001). **(d)** Bipartite network of predicted prokaryotic host genera (left) and associated viral genera or putative viral genus clusters (right). Node colours indicate host ecological group or viral taxonomic assignment. Edge width shows cumulative viral abundance and edge transparency show number of samples supporting the associations. **(e)** Spearman correlations between the abundance of prokaryotic host genera and the summed abundance of vOTUs predicted to infect the same host genus. Cell colour indicates Spearman ρ, with red representing positive correlations and blue showing negative correlations. Asterisks indicate BH-adjusted q < 0.05.

Next, we assessed whether the predicted ecological associations were reflected in abundance covariation between hosts and the collective vOTUs assigned to them. Predominantly positive virus-host relationships were observed across the major predicted host genera (Fig. 4E). The strongest associations were identified for *Staphylococcus* (ρ = 0.725, *p* < 0.0001), *Corynebacterium* (ρ = 0.666, *p* < 0.0001), *Streptococcus* (ρ = 0.590, *p* < 0.0001), and *Aerococcus* (ρ = 0.524, *p* < 0.0001). *Streptococcus* and *Aerococcus* were also represented by multiple links within the network.

### The oral microbiome undergoes age-dependent functional remodelling of carbohydrate utilisation

To determine whether the observed taxonomic succession across age groups was reflected in microbial functional potential, we examined KEGG orthologue (KO) profiles. The Weaner 1 communities exhibited a broader and a more even functional repertoire compared to later stages (Fig. 5A). KO richness was significantly higher in Weaner 1 than all other groups (all *p* ≤ 0.05). Shannon diversity followed a similar trend, with Weaner 1 exceeding all subsequent stages (all *p* ≤ 0.0001), while no significant differences were observed among Weaner 2, Finisher 1, and Finisher 2. These findings suggest that despite exhibiting lower taxonomic diversity, the post-weaning microbiome is characterised by greater functional diversity. Despite this, functional composition continued to reorganise as animals aged. When accounting for farm effects, Aitchison distances between KO profiles showed significant differences by age (R*^2^*=0.181, *p* < 0.001; Fig. 5B), with farm itself contributing an additional 24.9% of observed variation. All pairwise comparisons between age groups were significant after correction for multiple testing, except between Weaner 2 and Finisher 1. The largest functional divergence was observed between Weaner 1 and Finisher 2 (R*^2^*=0.209, *p* < 0.001). Given the importance of carbohydrate metabolism to microbial adaptation to changing dietary substrates, we analysed CAZYme potential and predicted substrate profiles. Glycoside hydrolases and glycosyltransferases dominated CAZyme repertoire at every age, together accounting for 86-88% of annotated CAZyme abundance (Fig. 5C). While this overall class structure was conserved, predicted substrate composition underwent developmental changes. Plant-associated substrates represented the largest functional category across all stages (50.3– 58.4%), with an increase observed after Weaner 1 stage. In contrast, host and milk-associated substrates declined from 23.5% in Weaner 1 to about 20% in later stages (Fig. 5E). Polysaccharide utilisation loci (PUL) homology predictions provided further resolution of this transition by identifying substrate-specific changes (Fig. 5D). Starch was the most abundant predicted PUL substrate and increased substantially from Weaner 1 to Finisher 2, with significant differences observed between Weaner 1 and each subsequent stage (*p* < 0.0001). Predicted utilisation of fructan and lactose also increased at later stages, while arabinan and human milk oligosaccharide-like substrate predictions were more prevalent in Weaner 1 (Fig. 5D; Fig. S3).

**Fig. 5:**
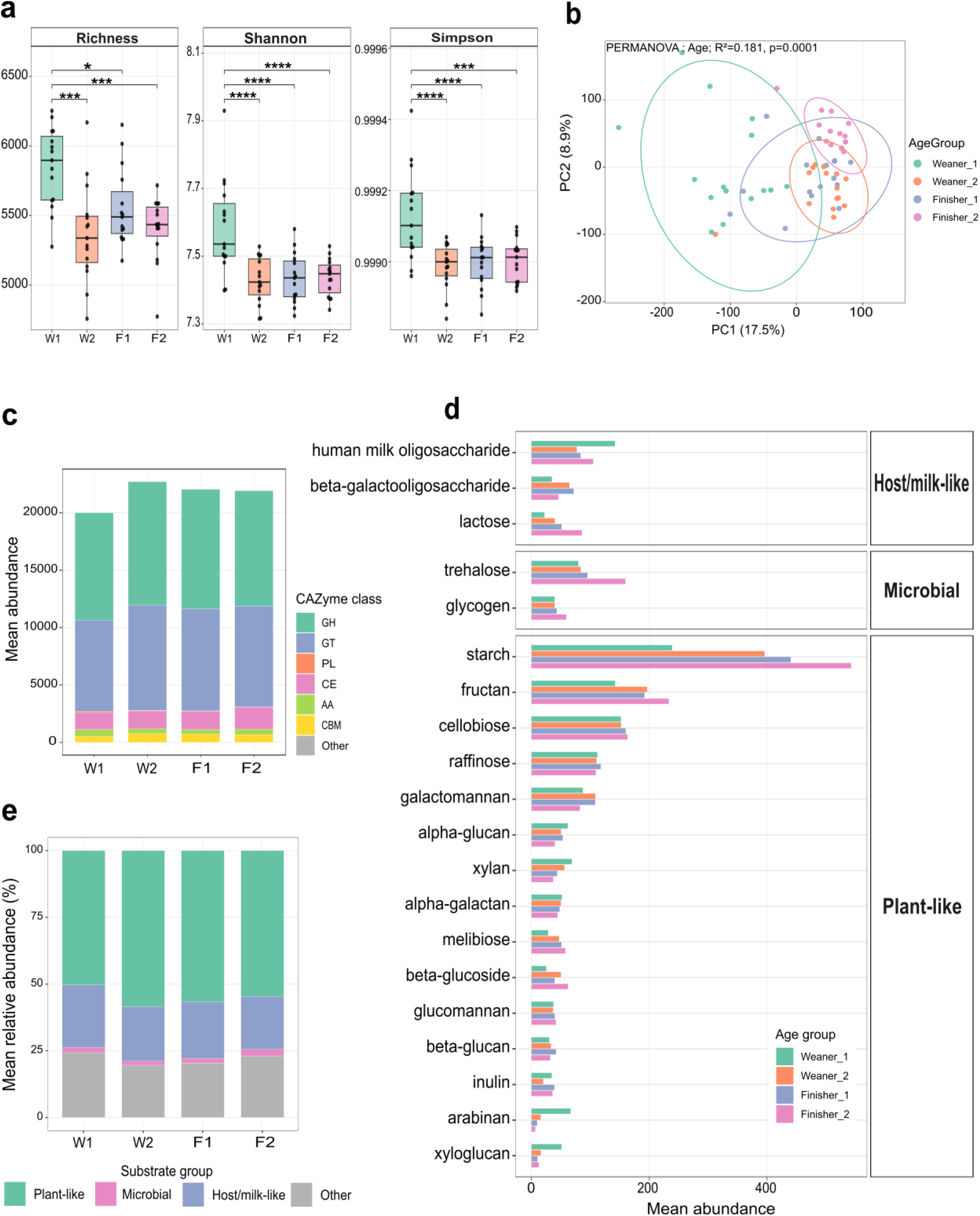
Variation in the functional potential of the oral microbiome across age groups. **(a)** Alpha diversity of KEGG orthologue (KO) profiles. **(b)** PCA of KO functional composition based on Aitchison distance from CLR-transformed abundance profiles. PERMANOVA adjusted for farm effects identified significant effect of age on functional composition (R^2^=0.18, p=0.0001). **(c)** Mean abundance of major CAZyme classes across age groups, including glycoside hydrolases (GH), glycosyltransferases (GT), polysaccharide lyases (PL), carbohydrate esterases (CE), auxiliary activities (AA), and carbohydrate binding modules (CBM). **(d)** Abundance of top predicted carbohydrate substrates across age groups. Bars show mean abundance for each substrate and are grouped by age. **(e)** Broader composition of predicted carbohydrate substrate groups across age groups. Bars show relative abundance of Plant-like, Most/milk-like, Microbial, and Other predicted substrates.

### Viral auxiliary functions partition by host association and predicted lifestyle

To assess whether the observed age-associated reshaping of the oral virome was reflected in its encoded functional potential, we analysed abundance-weighted viral gene categories. Phold improved functional resolution compared to Pharokka alone, with the proportion of genes classified as “Unknown” decreasing from 65.4% (abundance-weighted: 66.3%) with Pharokka to 41.0% (abundance-weighted: 42.5%) after Phold annotation. This improvement allowed for a more comprehensive depiction of the viral functional capacity. Although there was considerable taxonomic turnover across age groups, the overall functional architecture of the virome remained relatively stable (Fig. 6A). Among the resolved functions, head and packaging genes comprised 14.1–16.6%, DNA, RNA and nucleotide metabolism accounted for 12.9–13.5%, and tail-associated genes represented 10.9–12.7%. Structural functions, including head and packaging, tail, connector and lysis genes, became proportionally more prominent in Finisher 2, while auxiliary metabolic, moron and host-takeover genes remained stable at approximately 1.6–1.7%. In contrast, integration and excision genes were significantly abundant in Weaner 1 than in any other group (*p* < 0.0001). We subsequently examined the auxiliary and accessory gene pool that may influence virus-host interactions (Fig. 6B). Ranked by abundance-weighted support, toxin-antitoxin proteins were the most abundant, encoded by 367 vOTUs. Other abundant functions included phosphoadenosine phosphosulfate reductase (454 vOTUs), anti-CRISPR proteins (257 vOTUs), and NrdD-like anaerobic ribonucleotide reductases (424 vOTUs). Additional functions with high representation were PnuC-like nicotinamide mononucleotide transporters, glycosyltransferases, host nuclease inhibitors, restriction-alleviation proteins, and host-range or adsorption proteins. This suggests a broad capacity of the virome to modulate cellular metabolism, defence mechanisms, and host interactions.

**Fig. 6:**
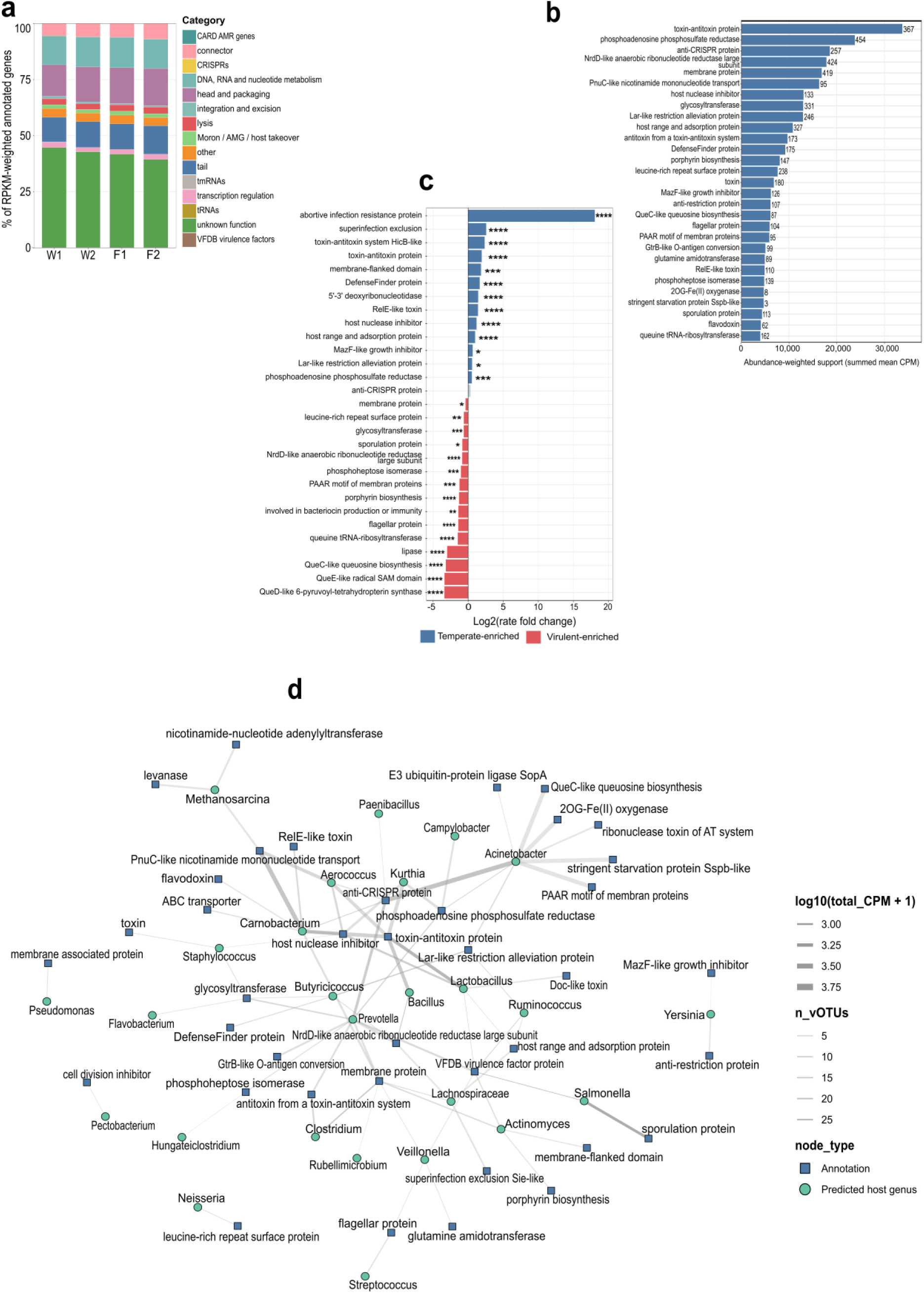
Functional profiles of the virome. **(a)** Abundance-weighted composition of predicted viral functional categories across age groups. Gene category counts for each vOTU were weighted by RPKM, and the stacked bars represent mean percentage contribution of each functional category **(b)** Top predicted accessory (moron, AMG, and host-takeover genes) functions ranked by abundance-weighted support. Bars represent summed mean abundance of vOTUs carrying each annotation, and the numbers next to the bar represent number of distinct vOTUs containing the annotation. **(c)** Lifestyle-specific enrichment patterns of moron, AMG and host-takeover genes. Log2 (fold-change) comparing temperate vs virulent phage rates for top annotations. Solid bars/colours indicate significant enrichment; transparent bars are non-significant. **(d)** Bipartite network linking predicted host genera to auxiliary/accessory annotations. Edge width reflects abundance-weighted support based on summed mean CPM and edge transparency reflects the number of distinct vOTUs supporting each host-annotation association.

To contextualise these functions within their predicted prokaryotic hosts, we mapped auxiliary and accessory genes to host genera inferred by PhaBOX2 (Fig. 6D). Viruses associated with *Acinetobacter* were enriched for anti-CRISPR proteins. PnuC-like nicotinamide mononucleotide transporters were prevalent among viruses predicted to infect *Carnobacterium* and *Aerococcus*. Toxin-antitoxin proteins were distributed across viruses linked to *Carnobacterium*, *Kurthia*, *Bacillus*, *Aerococcus*, and *Lactobacillus*, while host nuclease inhibitors were prominent in viruses associated with *Kurthia*, *Carnobacterium*, *Lactobacillus*, and *Aerococcus*. *Prevotella*-associated viruses contributed both anti-CRISPR and NrdD-like ribonucleotide reductase functions to the virome. These observations suggest that accessory functional potential is distributed in a non-random manner across predicted hosts rather than forming a uniform viral gene pool. Accessory functions also showed clear partitioning according to viral lifestyle (Fig. 6C). Toxin-antitoxin proteins were 3.83× more frequent in temperate phages compared to virulent phages (Log2FC: 1.94; *q* < 0.0001), and HicB-like toxin-antitoxin proteins were 5.11× more frequent in temperate phages (Log2FC: 2.35; *q* < 0.0001). Furthermore, DefenseFinder-associated proteins (3.13×; Log2FC: 1.65), RelE-like toxins (2.65×; Log2FC: 1.41), host-range or adsorption proteins (1.99×; Log2FC: 0.99), and host nuclease inhibitors (2.30×; Log2FC: 1.20) were all enriched (*q* < 0.0001) in temperate phages. Conversely, NrdD-like ribonucleotide reductases (1.81×; Log2FC: -0.85) were enriched in virulent phages. Also, queuosine-biosynthesis functions including QueD-like 6-pyruvoyl-tetrahydropterin synthase (10.51×), QueC-like proteins (9.12×), and QueE-like radical-SAM proteins (10.38×), were even more strongly enriched in virulent phages (all *q* < 0.0001). Thus, while the broad functional composition of the virome change only modestly with age, the improved resolution through structural homology (Phold) approach revealed a diverse accessory virome that is strongly structured by both predicted hosts and viral lifestyle.

### Age-associated shifts in resistome are linked to prokaryotic host turnover with limited viral contribution

We profiled the ARG landscape across age groups by normalising ARG abundance to RPKM. Weaner 1 had the highest ARG diversity, with a mean richness of 347±32 ARGs, which was significantly (*p* < 0.001) higher than Weaner 2 (282 ± 24), Finisher 1 (290 ± 26), and Finisher 2 (2280±18) (Fig. 7A). However, Shannon diversity remained stable across age groups. In addition to changes in ARG richness, the community composition of the resistome shifted significantly with age. PCA based on Aitchison distance revealed a separation of the prokaryotic communities ARGs by age group (Fig. 7B). PERMANOVA supported this observation, showing that age group accounted for a significant fraction of the variation in prokaryotic community (*R^2^*=0.145, *p* < 0.0001), with the strongest compositional pairwise difference observed between Weaner 1 and Finisher 2 (*R^2^*=0.174, *p* < 0.0001). Analysis of resistance mechanisms revealed that antibiotic efflux, antibiotic activation, target alteration, and target protection were predominant (Fig. 7C). In particular, in Weaner 1, efflux and inactivation mechanisms together constituted 66% of the resistome (both *p* < 0.0001), whereas the relative contribution declined to 49–54% in subsequent groups. A corresponding shift was observed in drug class composition in which lincosamide resistance was the most abundant in Weaner 1, together with macrolide, tetracycline, aminoglycoside and streptogramin resistance, while tetracycline resistance became dominant in subsequent ages (Fig. 7D).

**Fig. 7:**
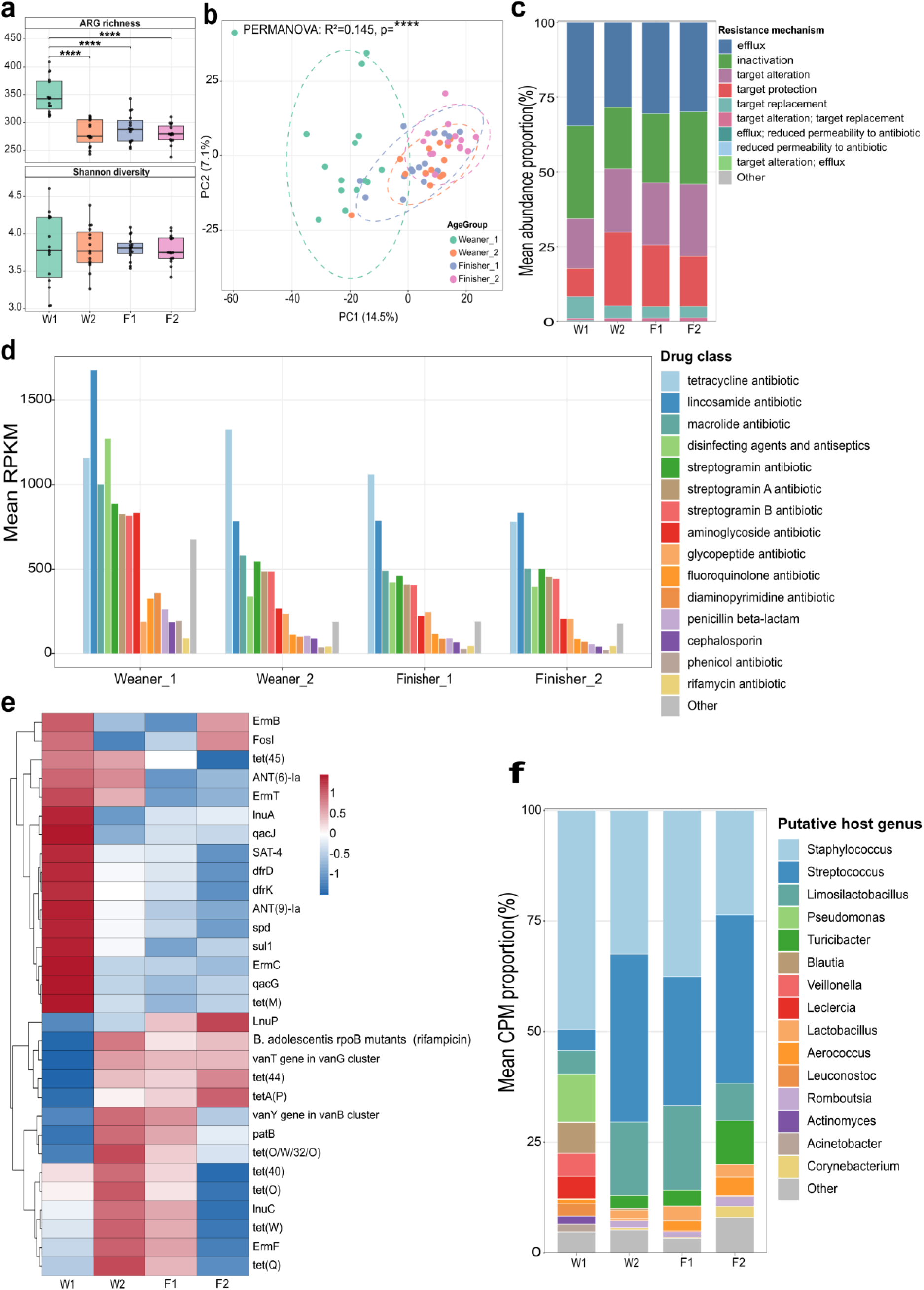
Patterns of antimicrobial resistance gene diversity and composition in pig oral microbiome. **(a)** Alpha diversity showing ARG richness (top) and Shannon diversity (bottom) across groups for prokaryotic communities. **(b)** PCA of prokaryotic resistome composition based on Aitchison distance from CLR-transformed ARO abundance profiles. Farm-adjusted PERMANOVA identified a significant age effect on resistome composition (R^2^=0.145, p < 0.001). **(c)** Mean relative abundance of the major resistance mechanisms across groups. **(d)** Mean abundance of drug classes across age groups based on RPKM from bulk microbiome libraries. **(e)** Heatmap of 30 most abundant genes across the groups. Colours indicate row-scales z-scores of log1p-transfoormed mean RPKM. Red represents high and blue shows lower abundance for a given ARG across age groups. **(f)** Relative composition of putative prokaryotic host genera carrying ARGs across groups.

At the antibiotic resistance ontology (ARO) level, Weaner 1 showed high abundance of *qacJ, lnuA, ErmB, ErmC, tet*(45), and *dfrK*, with several of these ARGs showing a substantial decline in abundance at later developmental stages (Fig. 7E). In contrast, abundance of *tet(W)* and *tet(O/W/32/O)* increased after Weaner 1 stage. These functional transitions were associated with significant turnover in the bacterial hosts associated with the ARGs (Fig. 7F). Of the ARGs that could be linked to a microbial host, *Staphylococcus* was the most dominant in Weaner 1, accounting for 49.5%, as well as *Pseudomonas* (10.8%), *Blautia* (7.0%), and *Limosilactobacillus* (5.3%). *Streptococcus* increased substantially in later groups, rising from 4.9% in Weaner 1 to 37.9% in Finisher 1, and 38.1% in Finisher 2. Other significant contributions from *Turicibacter*, *Aerococcus*, and *Corynebacterium* were observed at later ages.

In contrast to the prokaryotic resistome, the contribution of viruses was limited. Across the 25,223 vOTUs, only five ARGs were identified. These included *dfrG*, two *vanY* variants, *ErmY*, and *ANT(4’)-Ia*. After applying abundance and coverage thresholds, only three ARG-carrying vOTUs remained: a temperate vOTU encoding *dfrG* detected in 23.7% of samples; a virulent vOTU predicted to infect *Moraxella* and encoding *vanY/vanA*-cluster detected in 20.3% of samples; and a third vOTU encoding *vanY/vanB*-cluster vOTU detect in a single sample. These results indicate that representative oral vOTUs contributed minimally to the overall ARG reservoir compared to the prokaryotic community.

### *Streptococcus suis*-targeting phages are present in pig oral microbiome with age-associated abundance

To investigate the *S. suis*-associated component of the oral virome, we prioritised candidates identified independently by CHERRY host prediction and by nucleotide similarity to curated *S. suis*-associated INPHARED (pro)phages for ecological and comparative genomic analysis. This approach yielded 26 vOTUs, with a median length of 18.8 kb (10.4–37.7 kb). Representatives of all 26 vOTUs were temperate phages with only one identified as a provirus by geNomad. Recovery of these vOTUs was strongly influenced by both age and extraction method (Fig. 8A and 8B). The highest number of candidate vOTU was detected in Weaner 1 samples, especially in the bulk fraction, and richness declined at later production stages. In matched samples, bulk metagenomes consistently recovered more candidates than VLP libraries (*p* < 0.0001), with significant differences observed in Weaner 2, Finisher 1, and Finisher 2. Similarly, abundance was also affected by extraction method. Total abundance was consistently higher in bulk than their corresponding VLP enriched extractions (*p* < 0.0001). This observation contrasts with the higher overall viral richness recovered by VLP enrichment. Although VLP libraries were enriched approximately 11-fold for viral reads relative bulk (median 45.6% versus 4.5% of reads mapping to the 25,223 vOTUs), the lower abundance of *S. suis* vOTUs recorded in VLP indicates an approximately 100-fold over-representation of these sequences in the cellular fraction. Most individual candidates were detected sporadically across animals and farms (Fig. 8C). However, one vOTU, A21_NODE_293_length_25688, was present in 98.3% of samples and remained nearly ubiquitous across all production stages, with 100% prevalence in Weaner 1, Weaner 2, and Finisher 1, and 93.3% in Finisher 2. The next most prevalent candidate was found in only 20% of samples, and all other vOTUs occurred in 16.7% or fewer, giving a highly uneven prevalence structure dominated by a single widespread lineage.

**Fig. 8:**
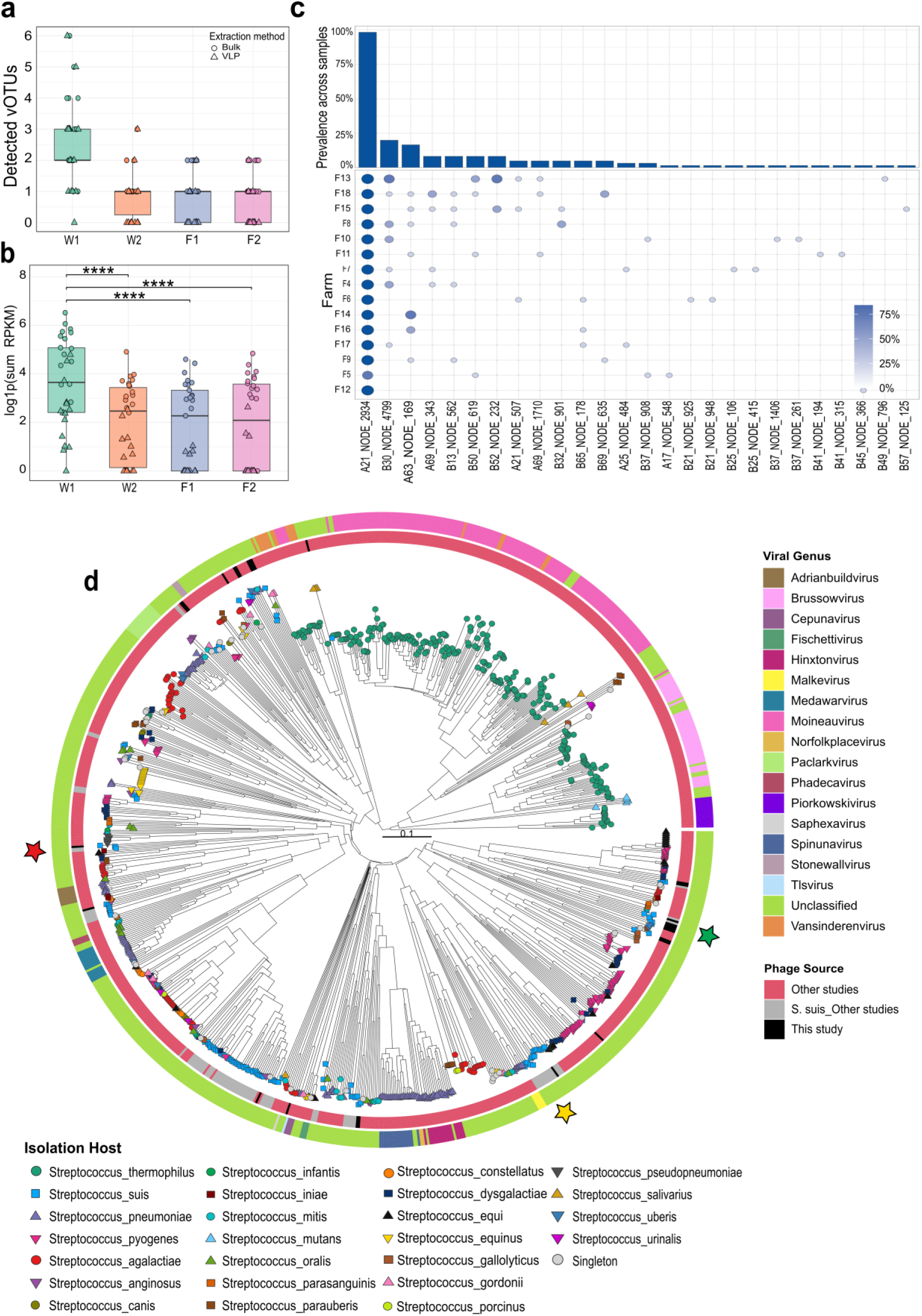
Recovery, abundance, and phylogenomic relationships of S. suis-associated vOTUs. **(a)** Number of S. suis vOTUs detected per sample across age groups. Points represent samples and shapes indicated extraction method (bulk or VLP). **(b)** Total abundance of S. suis-associated vOTUs across age groups. **(c)** Overall and farm-level prevalence of the 26 S. suis-associated vOTUs. Overall prevalence represents proportion of biological samples in which each vOTU was detected. Lower panel shows prevalence within individual farms. Point size and colour indicate farm-level prevalence. **(d)** Circular viral proteomic tree including the 26 S. suis vOTUs identified in this study and all streptococcal phage sequences in INPHARED database. Tip nodes are annotated by shape and colour to indicate the species of the host bacterium, with S. suis shown in a cyan-blue square, S. thermophilus in a green circle, and other streptococcal species in additional shapes and colours. The inner ring denotes the source of phages, including the S. suis vOTUs from this study (black), S. suis prophages from other studies (grey) and all other non-S. suis streptococcal phages from other studies (red). The outer ring represents the viral genus. Positions of cultivated S. suis phage Bonnie, Clyde, and SMP in the phylogeny are denoted by red, yellow, and green star, respectively.

Twenty-three of the 26 candidates were recovered from at least one size-filtered, DNase-treated VLP library. The three exceptions included A21_NODE_293_length_25688, which despite its near ubiquity in bulk fraction was not detected following VLP enrichment. The proportion of samples from which at least one VLP-recovered candidate could be identified declined significantly across production stages from 93.3% at Weaner 1 to 20.0% at Finisher 2. While *S. suis* was itself most abundant at Weaner 1, this did not account for the decline. Age group remained significant in a logistic model adjusted for *S. suis* relative abundance (likelihood ratio test *p* = 0.005). Likewise, sequencing effort did not cause the decline as depth did not differ across groups in either fraction (Kruskal-Wallis *p* ≥ 0.19). A reduction in candidate recovery was observed in both fractions between Weaner 1 and Finisher 2, however, this effect was more substantial following VLP enrichment, in which the mean number recovered per sample decreased from 2.4 to 0.3, compared to 2.9 to 1.3 in bulk fraction. Despite this decline, at least one candidate remained detectable in the bulk fraction for nearly all samples at each stage.

A viral proteomic tree showed that the 26 candidates did not form a single cohesive lineage (Fig. 8D). Instead, they formed part of multiple distinct clades predominantly constituted by other *S. suis* (pro)phages distributed across the broader streptococcal phage phylogeny. None of the *S. suis* phages, including the 26 candidates, could be assigned to a taxon at the genus or species level. The cultivated and characterised *S. suis* phages Bonnie (PQ720431), Clyde (PQ720432), and SMP (NC_008721) each occupied distinct positions within this diversity. This was supported by intergenomic similarity analysis which showed that none the the vOTUs were close species-level relatives of these three phages. The highest estimated intergenomic similarities to Bonnie, Clyde, and SMP were 13.0%, 40.9%, and 36.9%, respectively (Fig. S4). The most prevalent vOTU (A21_NODE_293_length_25688; 98.3% prevalence) was investigated further given its near-ubiquitous distribution across farms and age groups. This vOTU has a 25,688 bp genome encoding 40 predicted CDSs with complete structural modules including head/packaging, connector, and tail genes (Fig. S5). Functional annotation revealed a toxin-antitoxin system comprising a Doc-like toxin and cognate antitoxin, which may function as an addiction module to promote stable maintenance. The genome also encodes a DNA methyltransferase, potentially providing protection against host restriction-modification systems, alongside a suite of DNA replication genes including two helicases, a DNA polymerase, exonuclease, and ssDNA binding protein. BLASTN searches against the NCBI nucleotide database revealed high similarity (>90% nucleotide identity and >90% query coverage) to other *S. suis* prophages and presence in multiple *S. suis* genomes, with rare hits in closely related streptococci (*S. agalactiae* and *S. anginosus*), consistent with its identity as a predominantly *S. suis*-associated phage. The high sequence conservation across geographically distinct *S. suis* isolates, combined with its near-ubiquitous detection in pig oral samples and only sporadic occurrence in other streptococcal species, suggests this phage lineage is highly successful and well-adapted to *S. suis* as its primary host.

## Discussion

This study provides the first integrated metagenomic characterisation of the pig oral microbiome and DNA virome across four production stages. Consistent with previous reports, microbial diversity increased significantly after weaning. Early post-weaning pigs (Weaner 1) had lower species richness than older pigs, which indicates a rapid colonisation and diversification of the oral microbiome following the weaning transition [45]. However, this rise in richness did not translate to higher Shannon diversity, suggesting that while new taxa accumulate with age, community evenness remained constrained by a few dominant genera. *Streptococcus* persisted through all production stages in agreement with studies identifying it as a core oral taxon and the oral cavity as a significant reservoir for *S. suis* along with other related species [9, 46]. Early post-weaning pigs were enriched in *Pseudomonas* and *Staphylococcus*, whereas mid-nursery to early finishing stages were dominated by lactic acid bacteria, particularly *Lactobacillus amylovorus* and *Streptococcus alactolyticus*. By the late finisher stage, the community had become more even, with *Clostridioides*, *Aerococcus* and *Corynebacterium* rising alongside *Streptococcus*, *Nosocomiicoccus*, and *Anaerococcus*. This transition towards anaerobic and mucosa-associated taxa is characteristic of maturing oral biofilms, in which early colonisers deplete local oxygen and allow anaerobes to establish [47, 48]. Our shotgun metagenomic data extend these observations, providing species-level resolution.

In parallel, the composition of the oral DNA virome shifted significantly with age. Viral richness and Shannon diversity were lowest in Weaner 1 and increased thereafter, reflecting progressive diversification as the oral ecosystem matured. Most detected DNA viruses belonged to *Caudoviricetes*, consistent with other pig virome studies that report a dominance of bacteriophages over eukaryotic viruses [5]. However, only about 20% of vOTUs could be assigned a known genus, with nearly 40% occurring as singletons, highlighting the limited coverage of pig oral viromes in existing reference databases. *Astrithrvirus* was the most abundant assigned genus and increased after Weaner 1, while other genera remained comparatively minor. Similar trends have been observed in the gut virome of Tibetan pigs and other commercial breeds where community diversity and structure shift across development [49, 50]. Importantly, combining bulk and VLP-derived metagenomes increased recovery beyond either approach alone. VLP enrichment improved detection of viral diversity, consistent with studies showing enhanced recovery of low-abundance and more complete viral genomes [15, 51], while bulk metagenomes recovered 540 vOTUs that were not detected by VLP enrichment. This complementarity was also observed in bee and soil virome, where total metagenomes tend to sample dominant and temperate phages [52, 53]. As viral accumulation had not reached saturation, it is probable that even combined approach underestimates the true diversity present.

We observed strong coupling between prokaryotic and viral diversity and community structure. Prokaryotic and viral richness were significantly correlated, and beta diversity and Procrustes analysis showed that pigs with similar prokaryotic assemblages tended to carry similar viral communities. Virus-host linkage modelling supported this relationship at the taxon level, with abundance covarying among several major predicted hosts. For instance, *Streptococcus* was associated with *Fischettivirus* and *Vansinderenvirus*, while *Aerococcus* was linked to putatively novel genus clusters. These associations could reflect either virus (phage)-mediated regulation of bacterial populations or, conversely, phage dynamics being driven by host availability—such that, as some prokaryotic host decline with age, their associated phages decline proportionally. Irrespective of the directionality, the data demonstrates that oral prokaryotic and viral communities develop in tandem, mirroring patterns reported in human and pig gut microbiomes [54, 55].

We found that the relationship between taxonomic and functional diversity was not directly proportional. Weaner 1 had the lowest species richness but also possessed the broadest and most even KEGG orthologue repertoire, suggesting that taxa accumulating with age increasingly contributed functions already present in the community rather than extending its overall metabolic capabilities. Such functional redundancies is a recognised feature in microbial communities where phylogenetically distinct organisms encode similar functional capacities [56]. Nonetheless, the communities remained functionally distinct, with composition diverging across age groups along the same trajectory observed in taxonomic profiles. These patterns were exemplified in carbohydrate metabolism potential, where the distribution of CAZyme classes remained broadly conserved at each age group, while predicted substrate utilisation shifted progressively from host and milk-associated glycans towards plant-derived carbohydrates. Starch became the dominant predicted substrate as the diet transitioned from sow milk to cereal-based feed, trending the dietary transition that occurs during weaning in the pig gut [57]. Given that dietary substrates are encountered in the oral cavity prior to reaching the gastrointestinal tract, it is likely that the oral microbiota responds directly to dietary transitions and not passively reflecting downstream intestinal processes.

In contrast to the microbial community, the structural and nucleotide metabolism categories that dominate the virome showed little sensitivity to age. This is consistent with the conserved requirements for replication, DNA metabolism, virion assembly, and host lysis across tailed phage lineages. Integration and excision were the exception, being twice as abundant in Weaner 1 than other groups. Because these genes mediate prophage integration and excision, their enrichment suggests elevated lysogenic activity during early post-weaning. Structural homology searches with Phold reduced the proportion of unclassified genes from approximately two-thirds to 41%, supporting previous reports of improved annotation using structural approaches [35]. Nevertheless, the unassigned fraction was substantial, highlighting the persistence of functional dark matter within this niche. Beyond this otherwise broadly conserved functions, the accessory gene repertoire was more broadly structured by lifestyle. Temperate phages were enriched for toxin-antitoxin systems, defence-associated proteins, and host nuclease inhibitors. These functions have been implicated in antiphage defence and in the exclusion of secondary infecting phages from the lysogenised cell [58, 59]. Conversely, virulent phages harboured anaerobic ribonucleotide reductases—consistent with the nucleotide demands of productive lytic replication—as well as significant enrichment of queueosine biosynthesis genes. QueC/E/D are linked to protection against host endonucleases [60, 61], although distinguishing genes that augment host metabolic processes from those essential to viral processes remain challenging. Under a recently proposed auxiliary viral genes (AVG) framework, QueC/D/E likely represent essential genes for tRNA maturation and genome modification that promotes translation efficiency [7]. Accessory functions were also enriched according to host taxonomy. This pattern was expected, particularly for anti-CRISPR as these proteins inhibit specific Cas subtypes and must align with CRISPR-Cas systems present in their hosts [62]. Also, PnuC-like nicotinamide mononucleotide transporters were concentrated among viruses infecting *Carnobacterium* and *Aerococcus*, which is significant given that several bacterial defence systems function by depleting NAD+ and that phages have evolved pathways to restore it [63]. Overall, accessory gene content appears to be shaped by defensive strategies of the host as well viral replication strategy.

Antimicrobial resistome analysis demonstrated clear age-related patterns. ARG diversity and abundance were highest in Weaner 1 in agreement with established patterns in pig gut studies, where ARGs peak in young animals and decline during development [57, 64]. This may partly reflect the ecological replacement of antimicrobial-resistant early colonisers by more specialised commensal taxa during microbiome maturation. We identified that *Staphylococcus* harboured almost half of all host-linked ARGs in Weaner 1 before being mostly replaced by *Streptococcus* in finisher groups. A similar turnover was found at the gene level where *lnu(A), tet(45),* and *dfrK*, all characteristic of livestock-associated staphylococci [65], declined after Weaner 1, while *tet(W)*, and *tet(O/W/32/O)* increased. These ribosomal protection genes are common among commensal streptococci, oral, and gut anaerobes, [66, 67], linking resistome succession directly to changes in bacterial community structure. Although antimicrobial exposure contributes to emergence and spread of ARGs, previous studies have shown high abundance of ARGs in antibiotic-naïve pigs [64]. Thus, the elevated ARGs in younger pigs might be due to the persistence of maternal-derived resistant bacteria combined with environmental acquisitions during early post-weaning period, when microbiome instability creates opportunities for opportunistic colonisation. The high abundance of *qacJ* further supports the contribution of environmental acquisitions, as this gene confers resistance to quaternary ammonium disinfectants rather than to antibiotics [68]. Importantly, ARGs were overwhelmingly prokaryotic in origin, with only three detected in vOTUs after applying abundance and coverage thresholds. As mobile genetic elements, phages have been recognised as a reservoir for ARGs, however, our study and several previous reports demonstrate that their role might be overstated relative to other MGEs [17, 69, 70].

The currently available collection of cultivated *S. suis* phages remains very limited despite the recognised significance of the pathogen and extensive prophage prevalence in its genomes. Phage SMP was the only genome-characterised infective *S. suis* phage for many years until our recent isolation of Bonnie and Clyde revealed additional, genomically distinct lineages [23, 24]. The present analysis therefore asked whether the pig oral cavity—the major natural reservoir of *S. suis*—contains a broader phage community than has been captured through culture-based approaches. The candidate 26 vOTUs identified here were distributed across multiple streptococcal phage clades and exhibited minimal intergenomic similarity to SMP, Bonnie, or Clyde, with none reaching species-level similarity to the curated *S. suis* phages in INPHARED collection. Together with the extensive prophage diversity we previously identified in Irish *S. suis* genomes [25], these findings indicate that the cultivated phages represent only a small fraction of the phage diversity associated with this species. Importantly, only one of the 26 vOTUs was annotated as a provirus, while 23 were recovered from at least one size-filtered, DNase-treated VLP library, supporting their association with extracellular viral particles across most lineages, although this does not demonstrate infectivity. The mean number of candidates recovered between bulk and VLP fractions were similar at Weaner 1, but this similarity decreased over time. Given that this pattern persisted after adjusting for *S. suis* abundance and sequencing depth, it suggests that viral particle-associated recovery is structured by developmental stage and may reflect age-related differences in productive phage release, extracellular persistence, or host-phage interactions. Induction of temperate phages is known to respond to host physiology, microbial interactions, and environmental conditions [71, 72]. Moreover, the parallel enrichment of viral integration and excision functions at this stage further supports a heightened lysogenic turnover in early post-weaning pigs.

One vOTU displayed a distinct pattern. It occurred in 98.3% of samples and shared very high similarity with prophage sequences within *S. suis* genomes from diverse geographic regions. However, it was detected in only bulk but was not classified as provirus by geNomad, indicating a highly prevalent lineage associated with *S. suis* without evidence of viral particle release. Notwithstanding, the presence of several head and packaging, connector, and tail genes suggests that this lineage retains the machinery required for virion assembly. Moreover, the identification of a Doc-like toxin-antitoxin system and DNA methyltransferase suggests potential mechanisms for the persistence and evasion of host defences, especially considering the prevalence of restriction-modification systems in *S. suis* [25]. Overall, these findings expand the known diversity of *S. suis*-associated phages and establish a practical framework for phage discovery. Early post-weaning stage where bulk and VLP recovery were similar, may provide the best opportunity for direct isolation of extracellular candidates. In contrast, persistent bulk-associated lineages, such as A21_NODE_293_length_25688, may require induction or host-linked long-read sequencing approaches to establish whether they persist in integrated or other intracellular forms and whether they retain the capacity to exit these states and produce viral particles. Beyond intact phages, this expanded *S. suis*-associated vOTU catalogue may also provide a reservoir of phage-encoded proteins with therapeutic potential. For example, several *S. suis* prophage-derived lysins including LySMP, Csl2, Ly7917, PlySs2 and PlySs9, as well as holins such as HolSMP, have demonstrated potent bactericidal activity *in vitro* and *in vivo* against *S. suis* and other pathogens [73–77].

In conclusion, this study provides the first integrated characterisation of the pig oral microbiome and DNA virome, showing that both components undergo predictable, coordinated development across age groups. Early post-weaning was observed as a critical window characterised by high resistome diversity and high abundance of *S. suis* phage, representing an optimal target for microbiome-based interventions. By combining bulk and VLP sequencing, we establish a complementary approach for comprehensive viral characterisation that could underpin virome-informed phage discovery and biocontrol. Extending this framework to include RNA viruses will be essential, as these may capture clinically relevant pathogens and interactions undetected by DNA virome studies. Given that oral fluids are already widely used in herd monitoring, incorporating virome profiling alongside microbiome analysis offers a route toward precision livestock management. As the threat of antimicrobial resistance continues, virus-bacteria interaction networks may provide novel targets for sustainable interventions in animal agriculture.

## Data availability

The sequencing data generated in this study have been deposited in the NCBI Sequence Read Archive and are available under BioProject PRJNA1337625. Nucleotide sequence of the 26 vOTUs predicted to infect *S. suis* are available on Zenodo at https://doi.org/10.5281/zenodo.24424931

## Acknowledgements

This project (Improved Pig Health through the Novel Application of SynBio in Phage Therapy, 2020US-IRL201) was funded by the Irish Department of Agriculture, Food and the Marine, through the 2020 US-IRL R and D Partnership Call.

## Author contributions

**Emmanuel Kuffour Osei:** conceptualisation, data curation, formal analysis, investigation, methodology, visualisation, writing – original draft, writing – review and editing. **Mario Andre S Ornelas:** data curation and investigation. **Edgar Garcia Manzanilla:** funding acquisition, methodology, writing – review and editing. **Jennifer Mahony:** conceptualisation, funding acquisition and project management, investigation, methodology, writing – review and editing. **John G Kenny:** conceptualisation, funding acquisition and project management, investigation, methodology, writing – review and editing. All authors reviewed and approved the final manuscript.

